# Optimization of metagenomic detection method for human breast milk microbiome

**DOI:** 10.1101/2024.12.09.627599

**Authors:** Qiao Zhang, Yi Zhang, Jianjiang Zhu, Yajun Gao, Wen Zeng, Hong Qi

## Abstract

This study aims to optimize the metagenomic detection methodology of the human breast milk microbiome and analyze its composition. Twenty-two milk samples were collected from the left and right sides of lactating women during re-examinations at the Haidian Maternal and Child Health Hospital, Beijing. Microbial cell wall disruption parameters were optimized, and a nucleic acid extraction method was developed to construct a microbial DNA/RNA library. Metagenomic next-generation sequencing (mNGS) sequencing was performed, and microbial composition was analyzed using the k-mer Lowest Common Ancestor (LCA) method with a self-generated database constructed via Kraken2 software. Data showed Q20 > 95% and Q30 > 90%, with an average total data volume of 5,567 ± 376.6 Mb and non-human sequence data of 445.1 ± 63.75 Mb, significantly enhancing sequencing efficiency. The microbiome included 21 phyla, 234 genera, and 487 species, with Firmicutes and Proteobacteria as dominant phyla. Predominant genera included *Staphylococcus* and *Streptococcus*, and major species were *Staphylococcus aureus*, *Streptococcus bradystis*, and *Staphylococcus epidermidis*. Species levels exhibited significant variations among different individuals. Microbial profiles of left- and right-sided milk samples were consistent at the phylum, genus, and species levels. In addition to common bacteria, diverse viral, eukaryotic, and archaeal sequences were detected. This study refined metagenomic detection methods for human breast milk microbiota. Specific flora colonization occurred in healthy breast milk, with the left and right sides exhibiting both correlations and distinct flora environments.

**Importance:** Breast milk is a vital source of nutrition and immunity for infants, with its microbial composition playing a critical role in shaping the neonatal gut microbiome and supporting early development. However, technical challenges in detecting microorganisms in milk’s complex, lipid-rich environment have limited understanding of the diversity and function of these microbial communities. This study developed an optimized metagenomic sequencing method to analyze the microbial communities in breast milk from healthy mothers, identifying a wide array of bacteria, viruses, eukaryotes, and archaea. Key bacterial genera such as *Staphylococcus* and *Streptococcus* were predominant, with specific flora exhibiting inter-individual variability. Additionally, the study revealed distinct yet correlated microbial environments in the milk from the left and right breasts. These findings advance the understanding of breast milk microbiota and provide a foundation for exploring its implications for maternal and infant health.

## 1. Introduction

Breast milk microbiota and its metabolites contribute positively to the development of gastrointestinal microbiota, regulating host gene expression and supporting immune development in infants. Breastfeeding is a critical determinant of the composition and functionality of the gastrointestinal microbiome of an infant (1). Understanding the human milk microbiome and implementing targeted beneficial interventions for developing infants to optimize the microbiome in breast milk, donor milk, or formula can improve maternal and infant health throughout their lifespan. Additionally, investigating breast milk microbiota contributes to advancements in diagnosing infectious diseases with greater accuracy and efficiency(2, 3).

Microbiome research uses several techniques, including bacterial culture, amplicon sequencing, and mNGS (4, 5). Culture-based methods typically rely on specific nutrient media and environmental conditions to cultivate microorganisms, allowing direct observation and analysis of their morphological, physiological, and biochemical characteristics. However, this approach is limited because only a portion of culturable microorganisms can be detected and does not capture the full diversity of the microbial community (6–9).

Amplicon sequencing uses second/third-generation sequencing platforms to sequence targeted PCR products, such as 16S rRNA (ribosomal RNA), 18S rRNA, ITS (internal transcribed spacers), or functional genes. This approach addresses the drawbacks of traditional culturing by identifying non-culturable microorganisms and providing insights into the microbial community structure, evolutionary relationships, and their environmental correlations for environmental samples. It can directly detect the presence of microorganisms in the environment and is suitable for investigating rare or difficult-to-culture microorganisms. In recent years, 16S rRNA sequencing technology has been widely utilized to study the breast milk microbiome (5, 10). However, this method has certain limitations. Different hypervariable target regions of 16S rRNA can yield distinct phylum profiles from the same sample, and it is difficult to detect novel or highly variable microorganisms, such as viruses and fungi. Additionally, the limitation of the amplified region leads to the resolution problem of 16S rRNA at the microbial genus and species level (11).

mNGS is a high-throughput detection technique that sequences all nucleic acids within a specimen and employs bioinformatics analysis to identify microorganisms. Unlike traditional microbial nucleic acid diagnostic methods, mNGS does not require prior amplification of specific nucleic acids targeting the genome of one or more microorganisms. Instead, it efficiently and accurately captures the complete genomic information of a sample, enabling an unbiased analysis covering a wide range of microorganisms, including bacteria, fungi, viruses, parasites, mycoplasmas, and chlamydia (6, 12). Despite its potential, the application of mNGS in studying healthy milk microorganisms remains limited. Comprehensive analyses of viruses, eukaryotes, archaea, and other microorganisms in milk are scarce (13, 14).

One of the key challenges is that considerable fat, protein, and protease content in breast milk interfere with microbial detection, reducing the efficiency of microbial extraction and analysis. Consequently, developing effective nucleic acid extraction methods specific to milk is critical to facilitating the application of mNGS in breast milk research. Moreover, it is of considerable importance to develop a fast, sensitive, and effective pretreatment method suitable for clinical detection and to support the application of mNGS in the study of milk microorganisms.

This study introduces a novel human milk sample pretreatment method combined with metagenomic sequencing technology. Using this approach, breast milk microorganisms, including bacteria, viruses, fungi, and archaea, were detected and analyzed in detail at the phylum, genus, and species levels in the milk of healthy mothers.

## 2 Results

### 2.1 Optimization of Human Milk Metagenomic Sequencing Methodology

#### 2.1.1 Sample Pretreatment

The mean and standard deviation of the two CT values were analyzed in relation to the wall-breaking method used for pre-treated samples. Detailed CT data are provided in the supplemental material (Table s1).

When compared with the single pickling glass tube, the CT and mean values for the ternary wall-breaking tube were lower. The outcomes of the treatment employing a ternary wall-breaking tube were examined. Scheme 5 achieved a CT value of 28.7 ± 5.8, representing the lowest average value and low dispersion, as provided in Table 1.

**Table 1.** Comparison of CT values of wall breakage of pretreated milk samples from healthy lactating women.

| Option | 1 | 2 | 3 | 4 | 5 | 6 | 7 | 8 |
| --- | --- | --- | --- | --- | --- | --- | --- | --- |
| M-mean | 29.5 | 29.2 | 29.2 | 29 | 28.7 | 28.8 | 29.6 | 28.7 |
| M-SD | 5.5 | 5.7 | 5.8 | 5.9 | 5.8 | 6.1 | 6.3 | 6.1 |
| V-mean | 30.2 | 29.9 | 29.7 | 29.7 | 29.2 | 28.9 | 28.9 | 29 |
| V-SD | 5.7 | 5.8 | 6.2 | 5.9 | 6.4 | 6.4 | 6.5 | 6.7 |

M: M tube; V: V tube

Based on these results, subsequent experiments were conducted using the wall- breaking method of the ternary wall-breaking tube, as described in Scheme 5.

#### 2.1.2 Off-board Data Quality Control

Twenty-two milk samples from 11 healthy women were subjected to nucleic acid extraction, library construction, and sequencing using bridging PCR. After removing human sequences, the metagenomic sequences were obtained, classified, and summarized. The statistics for the effective sequence data are provided in Table 2. The quality metrics showed that Q20 > 95% and Q30 > 90%, meeting sequencing requirements and confirming the suitability of the specimens. The average total data size of the samples was 5,567 ± 376.6 Mb, while the average data size of non-human sequences was 445.1 ± 63.75 Mb. Details of the nucleic acid extraction concentrations of each sample are included in the supplemental material (Table s2).

**Table 2.** Summary of sequencing data for 11 samples.

M: M tube; V: V tube
| Participants | Raw data (Mb) | Raw reads (M) | Clean data (Mb) | Clean reads (M) | Human sequence (%) | Non-human reads (M) | Q20 (%) | Q30 (%) |
| --- | --- | --- | --- | --- | --- | --- | --- | --- |
| case 1L | 6,261.08 | 20.73 | 4,563.99 | 19.63 | 92.19 | 1.53 | 95.95 | 91.16 |
| case 1R | 4,140.86 | 13.71 | 3,014.97 | 13.13 | 92.86 | 0.94 | 95.1 | 90.34 |
| case 2L | 4,407.71 | 14.6 | 3,188.18 | 14.02 | 92.74 | 1.02 | 95.88 | 91.07 |
| case 2R | 3,830.95 | 12.69 | 2,820.88 | 12.24 | 93 | 0.86 | 95.83 | 91.03 |
| case 3L | 5,159.98 | 17.09 | 3,809.69 | 16.19 | 94.04 | 0.96 | 95.65 | 90.99 |
| case 3R | 5,478.17 | 18.14 | 3,820.06 | 16.9 | 93.67 | 1.07 | 95.36 | 90.37 |
| case 4L | 3,623.74 | 12 | 2,966.36 | 11.76 | 93.17 | 0.8 | 96.04 | 91.33 |
| case 4R | 3,717.34 | 12.31 | 2,909.81 | 11.85 | 92.48 | 0.89 | 96.35 | 91.79 |
| case 5L | 3,535.04 | 11.71 | 2,732.03 | 11.3 | 93.4 | 0.75 | 95.88 | 91.17 |
| case 5R | 3,516.54 | 11.64 | 2,761.44 | 11.37 | 93.73 | 0.71 | 95.45 | 90.62 |
| case 6L | 8,689.24 | 28.77 | 6,710.94 | 28.1 | 93.96 | 1.7 | 96.62 | 92.34 |
| case 6R | 8,656.4 | 28.66 | 6,268.79 | 27.43 | 93.82 | 1.69 | 96.02 | 91.59 |
| case 7L | 8,245.88 | 27.3 | 6,191.34 | 26.45 | 83.2 | 4.44 | 96.12 | 91.73 |
| case 7R | 7,130.66 | 23.61 | 5,322.86 | 22.84 | 83.52 | 3.76 | 95.99 | 91.38 |
| case 8L | 5,187.07 | 17.18 | 3,583.85 | 15.69 | 94.2 | 0.91 | 94.94 | 90.27 |
| case 8R | 5,793.87 | 19.19 | 4,271.79 | 18.66 | 94.84 | 0.96 | 96.17 | 91.57 |
| case 9L | 5,520.47 | 18.28 | 4,104.96 | 17.6 | 93.62 | 1.12 | 96.49 | 92.32 |
| case 9R | 4,184.34 | 13.86 | 3,198.82 | 13.5 | 86.87 | 1.77 | 96.05 | 91.67 |
| case 10L | 6,362.17 | 21.07 | 4,751.8 | 20.47 | 89.33 | 2.18 | 96.4 | 91.94 |
| case 10R | 8,858.41 | 29.33 | 6,631.19 | 28.5 | 94.84 | 1.47 | 96.55 | 92.07 |
| case 11L | 5,102.73 | 16.9 | 3,657.88 | 16.16 | 95.43 | 0.74 | 96.44 | 92.07 |
| case 11R | 5,078.06 | 16.81 | 3,571.94 | 15.7 | 94.24 | 0.9 | 95.68 | 91.21 |

### 2.2 Metagenome Sequencing of Healthy Maternal Breast Milk

#### 2.2.1. Species Distribution at Phylum, Genus, and Species Levels in Milk of Healthy Mothers

Microorganisms identified in the breast milk of healthy mothers were classified into 21 phyla, 234 genera, and 487 species. Analysis of the top 20 abundant phyla, genera, and species (Fig. 1) revealed that *Firmicutes* predominated at the phylum level, followed by *Proteobacteria*, which collectively accounted for more than 90% of the bacterial species.

**Figure 1.**
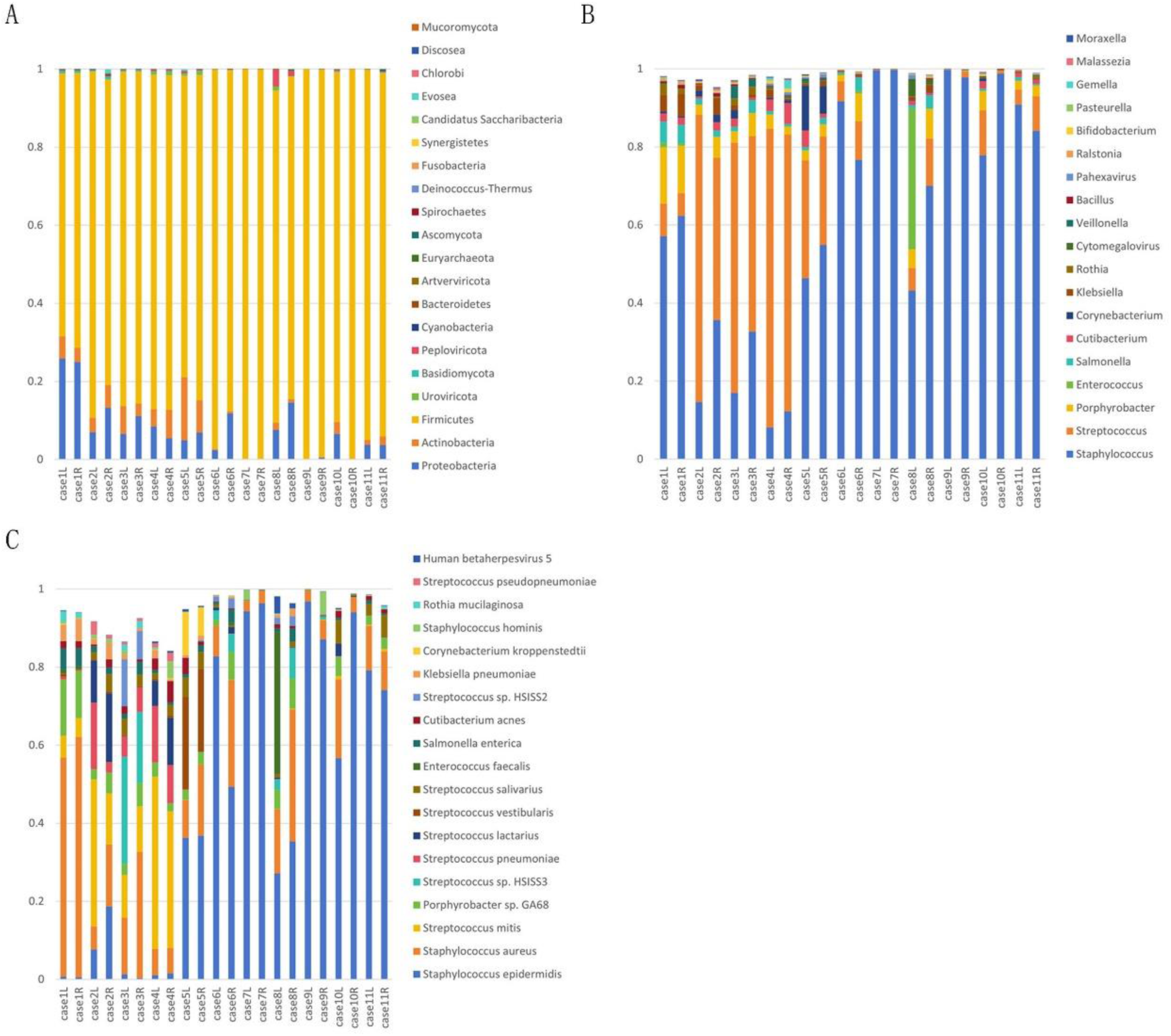
Microbial species distribution in the breast milk of healthy lactating women, depicted at the phylum, genus, and species levels. (A) Top 20 abundant phyla in the breast milk. (B) Top 20 abundant genera in the breast milk. (C) Top 20 abundant species in the breast milk.

In addition to common bacterial species, this study identified *Euryarchaeota* and detected fungi such as *Candidatus Saccharibacteria*, *Mucoromycota*, and *Planctomycetes*. Additionally, viruses like *Peploviricota*, *Uroviricota*, and *Artverviricota* were present.

At the genus level, *Staphylococcus* and *Streptococcus* constitute most of the microbiota. *Cytomegalovirus* and *Pahexavirus* virus were detected within the top 20 genera. Furthermore, *Malassezia* was prevalent in milk. *Bifidobacterium* was extensively present, while small quantities of *Lactobacillus* were observed in some individuals.

*S. epidermidis*, *S. aureus*, and *S. mitis*accounted for most of the microorganisms at the species level. The phylum and genus levels were largely consistent on both sides. However, significant variability was observed at the species level among different individuals.

#### 2.2.2. Alpha Diversity Index Analysis

Alpha diversity serves as a comprehensive indicator of community richness and evenness. The Chao1 index, which measures community richness, and the Shannon Index, which assesses both community richness and uniformity, were used to evaluate the diversity of microbial communities in milk (Table 3). The Chao1 index ranged from 59 to 216.6, while the Shannon index varied from 0.23 to 3.88. Alpha diversity showed fluctuations across samples. The number and abundance of species on the left and right sides of Case 3 milk exhibited significant dissimilarity. In contrast, the abundance and evenness of microbial communities were consistent between the left and right sides of the other samples.

**Table 3.** Alpha diversity indices for microbial communities in breast milk samples from 11 healthy mothers.

| Participants | Chao1 Index | Species Number | Shannon Index |
| --- | --- | --- | --- |
| case 1L | 86.4 | 67 | 2.55 |
| case 1R | 59 | 52 | 2.29 |
| case 2L | 159.5 | 140 | 3.33 |
| case 2R | 140.6 | 120 | 3.84 |
| case 3L | 183.2 | 160 | 3.88 |
| case 3R | 81 | 76 | 3.36 |
| case 4L | 164.2 | 128 | 3.35 |
| case 4R | 216.6 | 150 | 3.78 |
| case 5L | 93.3 | 80 | 2.87 |
| case 5R | 86.7 | 64 | 2.75 |
| case 6L | 85.7 | 76 | 1.16 |
| case 6R | 72.4 | 61 | 2.24 |
| case 7L | 84.6 | 80 | 0.41 |
| case 7R | 138.8 | 81 | 0.26 |
| case 8L | 80.2 | 71 | 2.63 |
| case 8R | 63 | 50 | 2.60 |
| case 9L | 82.1 | 65 | 0.23 |
| case 9R | 135.9 | 99 | 0.83 |
| case 10L | 168.3 | 111 | 2.27 |
| case 10R | 102.3 | 85 | 0.44 |
| case 11L | 69.1 | 59 | 1.25 |
| case 11R | 114.5 | 82 | 1.64 |

**Table 4.** Comparative analysis of metagenomic sequencing results from healthy breast milk samples.

**samples**
|  | Raw Data/Sample | Non-Human Sequences/Sample | Microbiome | Bacteria | Viruses | Eukaryotes | Archaea |
| --- | --- | --- | --- | --- | --- | --- | --- |
| Asnicar et al. (13) | 3.08Gb±1.5 | 26Mb±56 | - | not detailed | Caudovirales crAssphage | - | - |
| Ward et al.(15) | 1.33G(261.53 M*51bp/10=1 333.8Mb) | 1.33M*51bp/10=6. 78Mb | 360 prokaryotic genera | 360 genus | - | - | - |
| Our research | 5.56G(5567Mb±376.6) | 445.1Mb±63.75 | 21 phyla<br>234 genus<br>487 species | 12 phyla<br>194 genus<br>445 species | 3 phyla<br>13 genus<br>21 species | 5 phyla<br>14 genus<br>17 species | 1 phylum<br>4 genus<br>4 species |

#### 2.2.3 Comparative Analysis of Metagenomic Sequencing in Healthy Breast Milk

Compared with existing studies on the metagenomic analysis of microorganisms in healthy human breast milk using the Illumina sequencing platform, the average yield of our samples was 5,567 ± 376.6 Mb, of which 445.1 ± 63.75 Mb originated from non-human reads. This yield is considerably higher than the values reported in the other two metagenomic studies. Additionally, various viruses, eukaryotes, and archaea have been identified in breast milk. The sample pretreatment method markedly enhanced the sensitivity of Illumina metagenomic sequencing, enabling a more detailed classification of microorganisms at the phylum, genus and specie**s** levels (Table4).

#### 2.2.4 Viral, Eukaryotic, and Archaeal Sequences in Human Milk

Metagenomic sequencing of breast milk microorganisms identified sequences corresponding to viruses, eukaryotic microorganisms, and archaea. Phylum-level viruses include *Uroviricota*, *Peploviricota*, and *Artverviricota*, each containing diverse genera and species (Table 5). Eukaryotic microorganisms, such as fungi and protists, were detected. The principal phyla identified include *Basidiomycota*, *Ascomycota*, *Evosea*, *Discosea*, and *Mucoromycota*. Detailed classifications at the genus and species levels are provided in Table 6. Additionally, *Euryarchaeota* was detected in breast milk, encompassing several species at the genus and species levels (Table 7).

**Table 5.**
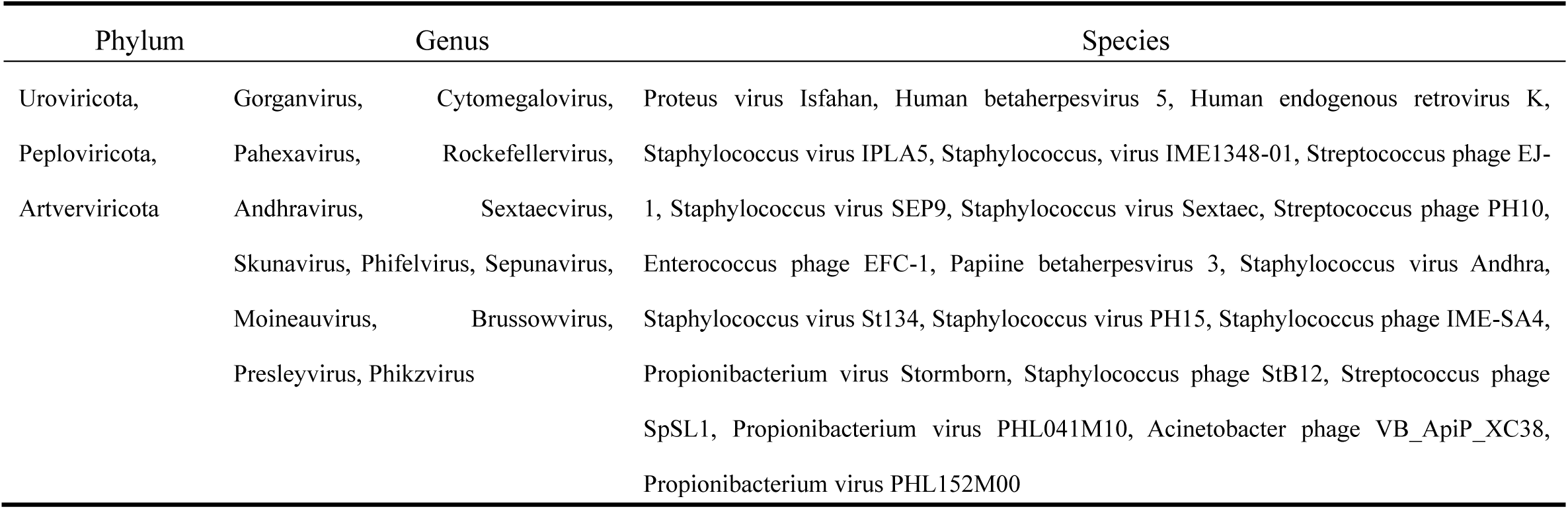
Viral Sequences Detected in the Human Milk Samples.

**Table 6.**
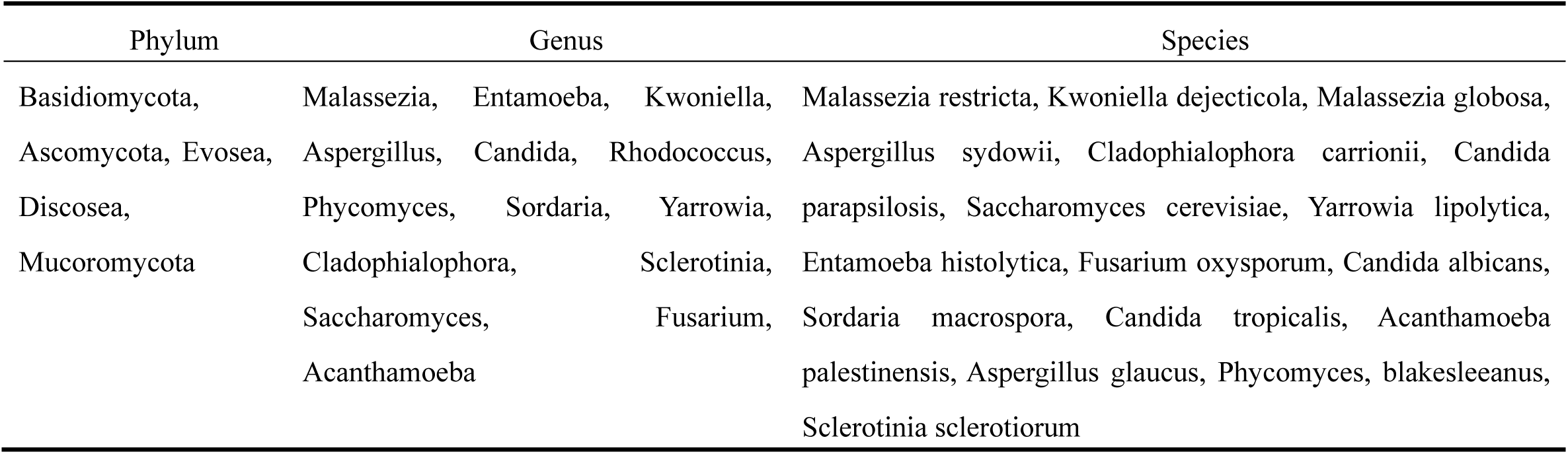
Eukaryotic Sequences in the Human Milk Samples.

**Table 7.**
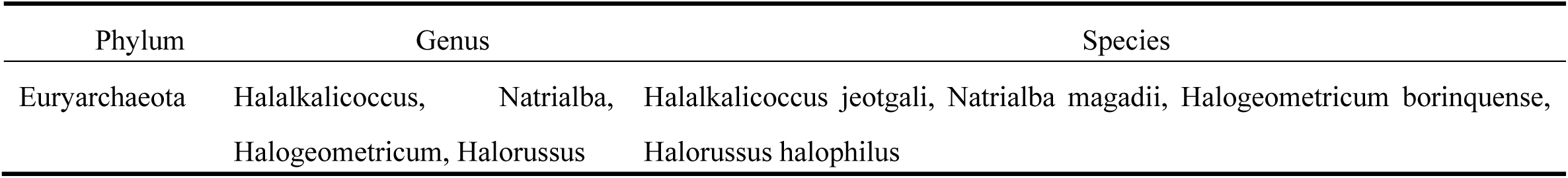
Archaeal Sequences in the Human Milk Samples.

## 3 Discussion

In this study, methodological optimization was performed, with gradient centrifugation as the primary approach to effectively remove a substantial quantity of milk fat and achieve stratified cell wall disruption. A ternary cell wall disruption tube, composed of glass beads, zirconium beads, and quartz sand, was used to enhance mechanical shear forces on complex microbial cell walls. This approach achieved a more comprehensive cell wall disruption effect than a single pickling glass cell wall disruption tube. Following optimization of the cell-wall disruption parameters, the extraction efficiency for fungi and certain bacteria with challenging cell wall to disrupt was markedly improved, facilitating the maximum extraction of complete microbial nucleic acids.

Building on these effective pretreatment methods, metagenomic sequencing technology was applied to detect and analyze milk microorganisms, including bacteria, archaea, eukaryotes, and viruses. Compared to the study by Ward et al. (15), which analyzed only prokaryotes in breast milk at the phylum and genus levels, this study extended microbial detection to the species level. Additionally, sequencing efficiency was markedly improved, and the average non-human sequence data in this study (445.1 Mb) far exceeded that reported by Asnicar et al. (26 Mb) (13). This study identified archaea, eukaryotes, and viruses in addition to common bacterial species in healthy breast milk, consistent with recent reports on milk microbial diversity (3).

At the phylum level, the types of bacteria identified were largely consistent with those reported in the literature; however, slight variations in composition were observed. *Proteobacteria* recorded the highest abundance in previous studies, while *Firmicutes* and *Proteobacteria* exhibited the highest abundance in the horizontal distribution, collectively accounting for more than 90% of the total bacterial species in healthy breast milk, with *Firmicutes* recording the highest abundance. These differences could be attributed to regional variations, sampling times, and differences in data analysis methods (11, 15).

In this study, *Staphylococcus* and *Streptococcus* were confirmed as the dominant genera at the genus level, consistent with findings from other non-culture-based methods, such as rRNA PCR, gas chromatography/mass spectrometry, and 16S rRNA sequencing (16, 17). In contrast, the proportion of *Pseudomonas* was relatively low in this study, with an average of only 0.09%. However, previous reports indicate that, in addition to *Staphylococcus* and *Streptococcus*, *Pseudomonas* are present in considerable numbers (11, 15). The variability in findings related to *Pseudomonas* remains controversial, with a few studies suggesting that *Pseudomonas* can be isolated from any sample, while others speculate that its high abundance may result from DNA extraction kits or other laboratory reagents (18, 19).

Species-level analysis revealed remarkable individual variation, likely related to maternal geographical location, diet, and lifestyle. Despite this variation, *S. epidermidis* and *S. aureus* were consistently identified as the predominant species in healthy breast milk. These findings align with previous species-level studies using both culture-based and non-culture-based methods (3, 5, 20).

In addition to common bacteria, this study identified various other microorganisms in breast milk. Virus including *Uroviricota*, *Peploviricota*, and *Artverviricota* were detected. Among fungi, *Basidiomycota* and *Ascomycota*, which have been reported previously, were detected along with *Discosea* and *Mucoromycota*, and an archaeon of the *Euryarchaeota*. Although individual viruses, fungi, and archaea were detected using other methods, our findings expand upon the limited and simplistic microbial classifications in earlier reports(2, 3).

Although many of the microorganisms identified belong to the same phylum as those of prior studies, considerable differences were observed at the genus and species levels (2, 3). Togo et al.(14) reported the presence of methanogenic archaea in breast milk. The distinction is that the Halobacteriaceae archaea identified in our study are consistent with those detected using 454 pyrosequencing; however, they represented distinct species categories (11). Additionally, our study is the first to report the presence of protozoan *Evosea* in breast milk.

Through microbial diversity analysis, the Chao1 index range in healthy maternal milk was determined to be 59–216.6, while the Shannon index ranged from 0.23 to 3.88. Differences in alpha diversity among individuals may be associated with variations in age, diet, geographic region, and work schedules of pregnant women. While the diversity of microflora in the left and right sides of breast milk of participants was generally similar, the number, richness, and evenness of the microflora were not entirely identical. In Case 3, the microflora diversity on the left side differed markedly from that on the right. This observation suggests that although the microbiota of the left and right sides is related, they may exhibit independent characteristics.

In summary, the pretreatment method established in this study markedly enhanced the detection capability of the metagenomic sequencing method. This approach enables the direct and effective extraction of DNA/RNA from all microbial genomes in breast milk for sequencing. Consequently, it facilitates a more comprehensive analysis of the diversity and functional characteristics of the microorganisms present in breast milk samples. Understanding the breast milk microbiome could have implications for maternal and infant health, particularly in areas like immunity, nutrition, or microbiota-related therapies. It frames the findings as a foundation for further research, which might explore specific interactions between microbiota and health outcomes or apply these methods to different populations.

## 4 Materials and Methods

### 4.1 Research Objects and Sample Collection

#### 4.1.1 Research Object

A total of 22 breast milk samples were collected from 11 breastfeeding women at the Haidian District Maternal and Child Health Hospital in 2023 at 42 days postpartum. All participants were healthy, with no signs of infection, such as fever, incision infections, mastitis, or endometritis, before and after delivery. The participants had no pregnancy complications, such as pregnancy-induced hypertension syndrome, severe anemia, or intrauterine growth restriction. None of the participants consumed antibiotics or probiotics within 1 month of delivery. All participants provided signed informed consent, which was approved by the local ethics committee.

#### 4.1.2 Milk Collection

Following iodophor disinfection, the initial breast milk was discarded. 5 mL of milk was collected from each side and promptly stored at -20℃ for other downstream analysis.

### 4.2 Methods

#### 4.2.1 Establishment of Sample Pretreatment Method

*Candida albicans* (Hebei Beina Biotechnology Co., Ltd., Hebei, China), *Streptococcus agalactiae* (Hebei Beina Biotechnology Co., Ltd., Hebei, China), and *Klebsiella pneumoniae* (Hebei Beina Biological Technology Co., Ltd., Hebei, China) were selected as representative fungi, gram-positive cocci, and gram-negative bacilli, respectively. These cultures were mixed, and a multiple glass bead, zirconium bead, and quartz sand ternary wall-breaking tube (M tube) were used and compared with a single pickling glass wall- breaking tube (V tube).

The parameters for the wall-breaking instrument, including oscillation speed (S), running time (T), interval time (D), and cycle number (C) were systematically adjusted. Nucleic acids were extracted and amplified using real-time quantitative PCR. Optimal wall- breaking parameters corresponding to the minimum and lowest CT value dispersions of the three microorganisms were selected by considering the mean and standard deviation (SD) of the CT value. In the wall-braking system, Bead Ruptor 12 (OMNI, Kennesaw, Georgia, USA) was used for pretreatment. Gradients of the wall-breaking parameters are listed in Table s3 in the supplemental material file.

#### 4.2.2 Nucleic Acid Extraction

A volume of 800 µL of milk sample was transferred to a 1.5 mL nuclease-free tube and centrifuged at 13,800×*g* for 5 min to separate the milk fat layer. The liquid below the cream layer was drained, and approximately 600 µL of the precipitate was transferred into a new 2.0 mL nuclease-free tube for use (where feasible to avoid the cream layer). A virus suspension was obtained through high-speed centrifugation, ensuring the integrity of viral genetic material by omitting wall-breaking treatment for viruses.

For samples requiring wall treatment, the process followed the optimized parameters earlier described. After the samples were pretreated, nucleic acids were extracted using the VAMNE Magnetic Pathogen DNA/RNA Kit (Vazyme Biotech Co., Ltd., nanjing, China) and quantified using Qubit 3.0(Invitrogen, Waltham, MA, USA).

#### 4.2.3 Paired-End Sequencing (PE) Library Construction and High-Throughput Sequencing

Microbial DNA/RNA libraries were constructed using 30 ng of nucleic acids, with the Rapid Max DNA Lib Prep Kit for Illumina (ABclonal, wuhan, China) used for library construction. After library construction, the concentration was measured using a Qubit 3.0 fluorescence quantifier, confirming a concentration exceeding 0.1 ng/µL. Metagenomic sequencing was performed on the Illumina NovaSeq6000 platform (Illumina, CA, USA), generating a data volume of 20 million reads.

### 4.3 Data Analysis

Bowtie2 was used to compare the original filtered sequence with the human reference genome (hg19). The human source sequences were excluded, and the k-mer LCA method was used to compare species with the self-owned data constructed using the Kraken2 software. A proprietary database was constructed using Kraken2 for species comparison. This database includes all bacterial, archaeal, fungal, protozoan, and viral species in the NCBI RefSeq database (Release 210, January 7, 2022), comprising approximately 19,400 species, including common pathogens. Data analysis was performed using QIIME Version 2:2020.2.0 software(21). Species abundance was calculated by RPM, (the number of organism reads per million sequence reads, Taxon Reads x 106 / Total Reads) represented by the formula: The total number of reads should be employed under appropriate circumstances to standardize the total sequence number of samples, which is conducive to inter-sample comparison and positive determination.

Statistical Methods: SPSS22.0 (IBM, Armonk, New York, USA) was used to analyze the mean and SD of parameters for statistical analysis.

## Acknowledgments

This work was funded by the High-level Talents Development Program of the Health System in Haidian District, Beijing, China (l2022HDXD004) and Hygiene and Health Development Scientific Research Fostering Plan of Haidian District Beijing (HP2024-30- 103002).

We would like to thank Editage (www.editage.cn) for English language editing.

---

**Table s1.**
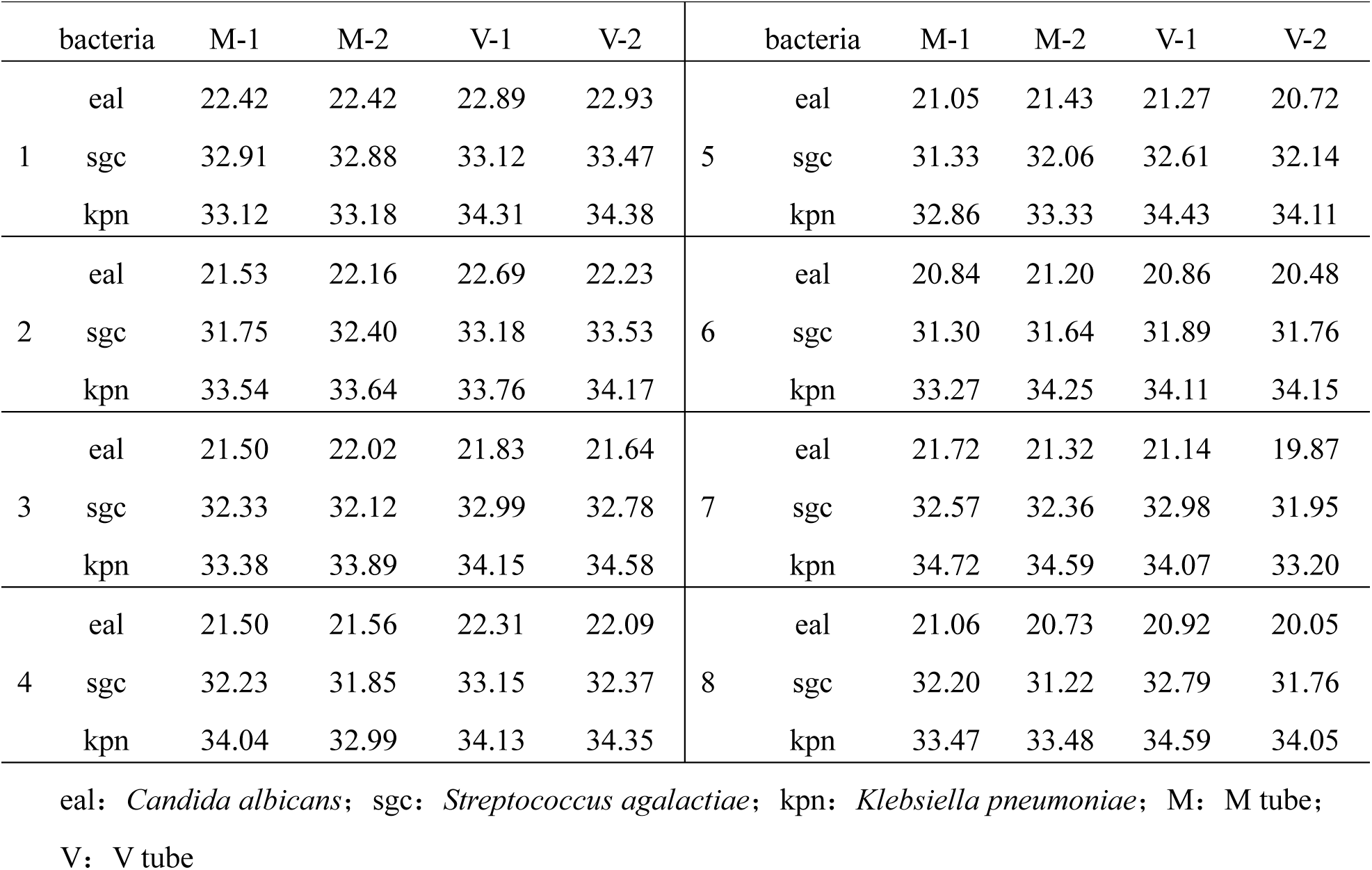
CT values of pretreatment samples.

**Table s2.** Extraction concentration of nucleic acid from 11 healthy breastfeeding women (ng/μl)

**Tabl s2 Extraction concentration of nucleic acid from 11 healthy breastfeeding women**
|  | (ng/μl) |  |  |  |  |  |  |  |  |  |  |
| --- | --- | --- | --- | --- | --- | --- | --- | --- | --- | --- | --- |
|  | case | case | case | case | case | case | case | case | case | case | case |
|  | 1 | 2 | 3 | 4 | 5 | 6 | 7 | 8 | 9 | 10 | 11 |
| left | 2.6 | 0.99 | 2.16 | 0.89 | 1.38 | 1.47 | 2.64 | 1.34 | 0.63 | 1.31 | 0.85 |
| right | 1.65 | 1.09 | 3.08 | 1.38 | 1.23 | 1.05 | 1.99 | 0.5 | 1 | 0.63 | 7.84 |

**Table s3.** Gradients of the wall-breaking parameters in pretreatment sample.

|  |  |  |  |  |  |  |  |  |
| --- | --- | --- | --- | --- | --- | --- | --- | --- |
| Parameter | Gradient |  |  |  |  |  |  |  |
|  | 1 | 2 | 3 | 4 | 5 | 6 | 7 | 8 |
| S (m/s) | 5 | 5 | 5 | 5 | 5.5 | 5.5 | 5.5 | 5.5 |
| T (s) | 40 | 50 | 40 | 50 | 40 | 50 | 40 | 50 |
| D (min) | 3 | 3 | 3 | 3 | 3 | 3 | 3 | 3 |
| C | 3 | 3 | 4 | 4 | 3 | 3 | 4 | 4 |

S: oscillation speed T: running time D: interval time C: cycle number

## References

1. Jost T, Lacroix C, Braegger C, Chassard C. Impact of human milk bacteria and oligosaccharides on neonatal gut microbiota establishment and gut health. Nutr Rev. 2015;73(7):426–37.

2. Stinson LF, Sindi ASM, Cheema AS, Lai CT, Mühlhäusler BS, Wlodek ME, Payne MS, Geddes DT. The human milk microbiome: who, what, when, where, why, and how? Nutr Rev. 2021;79(5):529–43.

3. Dombrowska-Pali A, Wiktorczyk-Kapischke N, Chrustek A, Olszewska-Słonina D, Gospodarek- Komkowska E, Socha MW. Human Milk Microbiome-A Review of Scientific Reports. Nutrients. 2024;16(10).

4. Miller RR, Montoya V, Gardy JL, Patrick DM, Tang P. Metagenomics for pathogen detection in public health. Genome Med. 2013;5(9):81.

5. Jost T, Lacroix C, Braegger C, Chassard C. Assessment of bacterial diversity in breast milk using culture-dependent and culture-independent approaches. Br J Nutr. 2013;110(7):1253–62.

6. Gu W, Miller S, Chiu CY. Clinical Metagenomic Next-Generation Sequencing for Pathogen Detection. Annual review of pathology. 2019;14:319–38.

7. Han D, Li Z, Li R, Tan P, Zhang R, Li J. mNGS in clinical microbiology laboratories: on the road to maturity. Critical reviews in microbiology. 2019;45(5-6):668–85.

8. Wilson MR, Naccache SN, Samayoa E, Biagtan M, Bashir H, Yu G, Salamat SM, Somasekar S, Federman S, Miller S, Sokolic R, Garabedian E, Candotti F, Buckley RH, Reed KD, Meyer TL, Seroogy CM, Galloway R, Henderson SL, Gern JE, DeRisi JL, Chiu CY. Actionable diagnosis of neuroleptospirosis by next-generation sequencing. The New England journal of medicine. 2014;370(25):2408–17.

9. Martín R, Jiménez E, Heilig H, Fernández L, Marín ML, Zoetendal EG, Rodríguez JM. Isolation of bifidobacteria from breast milk and assessment of the bifidobacterial population by PCR-denaturing gradient gel electrophoresis and quantitative real-time PCR. Applied and environmental microbiology. 2009;75(4):965–9.

10. Asenova A, Hristova H, Ivanova S, Miteva V, Zhivkova I, Stefanova K, Moncheva P, Nedeva T, Urshev Z, Marinova-Yordanova V, Georgieva T, Tzenova M, Russinova M, Borisova T, Donchev D, Hristova P, Rasheva I. Identification and Characterization of Human Breast Milk and Infant Fecal Cultivable Lactobacilli Isolated in Bulgaria: A Pilot Study. Microorganisms. 2024;12(9).

11. Jiménez E, de Andrés J, Manrique M, Pareja-Tobes P, Tobes R, Martínez-Blanch JF, Codoñer FM, Ramón D, Fernández L, Rodríguez JM. Metagenomic Analysis of Milk of Healthy and Mastitis-Suffering Women. J Hum Lact. 2015;31(3):406–15.

12. Chiu CY. Viral pathogen discovery. Current opinion in microbiology. 2013;16(4):468–78.

13. Asnicar F, Manara S, Zolfo M, Truong DT, Scholz M, Armanini F, Ferretti P, Gorfer V, Pedrotti A, Tett A, Segata N. Studying Vertical Microbiome Transmission from Mothers to Infants by Strain-Level Metagenomic Profiling. mSystems. 2017;2(1).

14. Togo AH, Grine G, Khelaifia S, des Robert C, Brevaut V, Caputo A, Baptiste E, Bonnet M, Levasseur A, Drancourt M, Million M, Raoult D. Culture of Methanogenic Archaea from Human Colostrum and Milk. Sci Rep. 2019;9(1):18653.

15. Ward TL, Hosid S, Ioshikhes I, Altosaar I. Human milk metagenome: a functional capacity analysis. BMC Microbiol. 2013;13:116.

16. Hunt KM, Foster JA, Forney LJ, Schütte UM, Beck DL, Abdo Z, Fox LK, Williams JE, McGuire MK, McGuire MA. Characterization of the diversity and temporal stability of bacterial communities in human milk. PloS one. 2011;6(6):e21313.

17. Fitzstevens JL, Smith KC, Hagadorn JI, Caimano MJ, Matson AP, Brownell EA. Systematic Review of the Human Milk Microbiota. Nutrition in clinical practice : official publication of the American Society for Parenteral and Enteral Nutrition. 2017;32(3):354–64.

18. Tanner MA, Goebel BM, Dojka MA, Pace NR. Specific ribosomal DNA sequences from diverse environmental settings correlate with experimental contaminants. Appl Environ Microbiol. 1998;64(8):3110–3.

19. Barton HA, Taylor NM, Lubbers BR, Pemberton AC. DNA extraction from low-biomass carbonate rock: an improved method with reduced contamination and the low-biomass contaminant database. J Microbiol Methods. 2006;66(1):21–31.

20. Fernández L, Langa S, Martín V, Maldonado A, Jiménez E, Martín R, Rodríguez JM. The human milk microbiota: origin and potential roles in health and disease. Pharmacol Res. 2013;69(1):1–10.

21. Bolyen E, Rideout JR, Dillon MR, Bokulich NA, Abnet CC, Al-Ghalith GA, Alexander H, Alm EJ, Arumugam M, Asnicar F, Bai Y, Bisanz JE, Bittinger K, Brejnrod A, Brislawn CJ, Brown CT, Callahan BJ, Caraballo-Rodríguez AM, Chase J, Cope EK, Da Silva R, Diener C, Dorrestein PC, Douglas GM, Durall DM, Duvallet C, Edwardson CF, Ernst M, Estaki M, Fouquier J, Gauglitz JM, Gibbons SM, Gibson DL, Gonzalez A, Gorlick K, Guo J, Hillmann B, Holmes S, Holste H, Huttenhower C, Huttley GA, Janssen S, Jarmusch AK, Jiang L, Kaehler BD, Kang KB, Keefe CR, Keim P, Kelley ST, Knights D, Koester I, Kosciolek T, Kreps J, Langille MGI, Lee J, Ley R, Liu YX, Loftfield E, Lozupone C, Maher M, Marotz C, Martin BD, McDonald D, McIver LJ, Melnik AV, Metcalf JL, Morgan SC, Morton JT, Naimey AT, Navas-Molina JA, Nothias LF, Orchanian SB, Pearson T, Peoples SL, Petras D, Preuss ML, Pruesse E, Rasmussen LB, Rivers A, Robeson MS, 2nd, Rosenthal P, Segata N, Shaffer M, Shiffer A, Sinha R, Song SJ, Spear JR, Swafford AD, Thompson LR, Torres PJ, Trinh P, Tripathi A, Turnbaugh PJ, Ul-Hasan S, van der Hooft JJJ, Vargas F, Vázquez-Baeza Y, Vogtmann E, von Hippel M, Walters W, Wan Y, Wang M, Warren J, Weber KC, Williamson CHD, Willis AD, Xu ZZ, Zaneveld JR, Zhang Y, Zhu Q, Knight R, Caporaso JG. Reproducible, interactive, scalable and extensible microbiome data science using QIIME 2. Nat Biotechnol. 2019;37(8):852–7.

